# Characterizing reward sensitivity to natural singing: an individual differences approach

**DOI:** 10.64898/2026.05.04.722621

**Authors:** Emma Segura, Urbano Lorenzo-Seva, Robert Zatorre, Boris A. Kleber, Antoni Rodríguez-Fornells

**Author notes:** **Corresponding author:** Antoni Rodríguez-Fornells, ICREA Research Professor, Department of Cognition, Development and Educational Psychology University of Barcelona, Campus Bellvitge, Feixa Llarga, s/n, 08907 L’Hospitalet de Llobregat, 08907 Barcelona, Spain. Both authors declare co-senior equal contribution (Boris A. Kleber, Antoni Rodríguez-Fornells).

## Abstract

Singing is an innate human behaviour present across cultures and the lifespan. Despite lacking direct biological advantages, its ubiquity suggests that it is intrinsically rewarding. This research aimed to investigate the underlying factors that explain variability in sensitivity to deriving reward and enjoyment from natural singing in the general population. In Study 1 (n = 606), an initial pool of items describing daily, non-professional singing behaviours were administered to an international adult sample. Exploratory factor analysis revealed a unidimensional structure of 20 items with acceptable model fit, organized into five facets representing distinct domains of singing-related rewards: 1) *pleasure and emotional evocation*, 2) *social singing reward*, 3) *singing frequency*, 4) *mood regulation through singing*, and 5) *inattentional singing during routine tasks*. In Study 2 (n = 430), confirmatory factor analysis in a new sample supported this structure. When both samples were combined (n = 1036), the unidimensional model defined by these five facets showed acceptable to excellent goodness-of-fit indices, supporting the conceptualization of singing reward as a multidimensional construct with differentiated facets. This led to the Barcelona-Aarhus Natural Singing Engagement Questionnaire (BANSEQ), which demonstrated excellent reliability (α = .94) and population-level stability. Study 3 (n = 1036) tested the convergent validity of BANSEQ with measures of music reward and engagement and identified sociodemographic and psychological correlates across the five facets of singing reward. Overall, these findings characterize the sources of individual differences in the hedonic experience of natural singing and propose BANSEQ as a robust psychometric tool for its assessment in the general population.

## Introduction

Singing, one of the most natural and direct forms of music-making, is an innate human behaviour that is ubiquitously present in daily life across diverse social contexts and cultures. For instance, parents sing lullabies to help infants sleep, teachers use songs to facilitate learning, relatives sing *Happy Birthday* to celebrate others, and people sing anthems to express group identity (e.g., a country or team) (Trehub & Trainor, 1998). Although singing does not provide any direct biological advantages, many individuals voluntarily engage in singing behaviours across their lifespan, either individually or in groups, to seek gratification, social interaction, or other socio-emotional benefits (Brown, 2000; Cohen, 2020; Mehr et al., 2019). Consistent with this view, prior research has demonstrated positive effects of singing on mental health in both clinical and non-clinical contexts. Individual and choir singing have been shown to enhance mood, positive affect, personal growth, interest in life, and emotional expression, as well as to increase social bonding and reduce stress and loneliness across healthy populations and individuals with psychiatric or neurological conditions (Fancourt et al., 2015; Insel & Fernald, 2004; Noice & Noice, 2008; Siponkoski et al., 2023; Smith et al., 2022; Stone et al., 2018; Weinstein et al., 2016; Williams et al., 2018). Beyond these psychological effects, singing has also been shown to increase respiratory and immune capacity in healthy populations (Kaasgaard et al., 2022; Kuhn, 2002; Thomasson & Sundberg, 1997), and to enhance language functions in individuals with acquired brain injury, Parkinson’s disease, and autism (Schlaug, G., Marchina, S., and Norton, 2008; Wan et al., 2010).

Although the evolutionary purpose of singing remains unclear, its ubiquity and central role in human life underscore its deep roots in our species (Mehr et al., 2019, 2021). One proposed mechanism underlying this persistence in human lineage across millennia is the close coupling of singing with brain reward networks, which may support the intrinsic motivation to engage in and repeatedly seek singing-related pleasurable experiences (Ferreri et al., 2019; Grewe et al., 2005; C. A. Wang et al., 2018). Neuroimaging studies have demonstrated that music listening engages brain regions involved in pleasure, emotion, and reward (Hodges, 2010). However, individuals vary substantially in the degree of pleasure they derive from music, reflecting large differences in how musical stimuli are perceived and experienced (Mas-Herrero et al., 2013). Such variability has been linked to individual differences in emotional sensitivity to music, the use of music for mood regulation, and enjoyment of music in social contexts, factors that are thought to mediate musical pleasure (Freeman, 2000; Juslin, 2003; Juslin & Västfjäll, 2008; Laeng et al., 2016).

In contrast to music listening, singing is an active, performative behaviour shaped by multiple physiological, psychological, cognitive, social, emotional, and contextual factors (Cohen et al., 2020; Juslin & Sloboda, 2012; Sloboda, 2000). Despite the complexity of singing and the potential variability in how it is experienced, most individuals in the general population report enjoying singing. In a sample of students without formal vocal training, 88.3% reported enjoying singing, primarily because it affords opportunities for self-expression, aesthetic experience, stress reduction, and mood regulation. However, about 12% of participants reported being reticent to sing, indicating that they enjoyed singing only when alone due to embarrassment about performing in front of others, or that they did not enjoy singing at all because dissatisfaction with their own voice elicited negative and self-critical thoughts (Chong, 2010). Further research has shown that, although individuals in the general population can sing with reasonable accuracy, both children and adults tend to misestimate their singing ability, which reduces their motivation to join in singing activities (Demorest et al., 2017). However, a large-scale study (n = 632) examining inherent singing skills across the lifespan (ages 6-99) and a wide range of musical experience (0-55 years of musical training) reported only a weak association between singing accuracy and singing enjoyment (*r* = .18; *p* < .01) (Pfordresher & Demorest, 2020). Together, these findings suggest that singing ability might not be a critical determinant of singing enjoyment, and that other factors likely influence singing engagement across the lifespan. This pattern converges with recent evidence indicating that individual differences in music reward sensitivity can be dissociated from overall musical aptitude. For example, Bignardi et al. (2024) demonstrated that musical aptitudes and musical enjoyment follow partly distinct neurobiological — and possibly genetic — pathways, suggesting that the capacity to derive pleasure from music-related activities may have evolved independently from musical skill. Consistent with this dissociation, neuropsychological evidence shows that music anhedonia (loss of pleasure for music listening) and amusia (loss of music perceptual abilities) can be selectively disrupted following brain damage (Mas-Herrero et al., 2014; Satoh et al., 2011). Overall, this body of research indicates that musical aptitudes — here operationalized as singing ability — do not necessarily predict greater sensitivity to reward and pleasure in music-related activities among the general population.

Beyond singing ability and enjoyment, relatively little is known about variability in sensitivity to natural singing reward among the general population. Natural singing can be defined here as any singing behaviour that emerges early in life and is maintained across the lifespan for sociocultural purposes or intrinsic enjoyment, independent of professional training or skill mastery. Within this framework, the present research aimed to identify and examine the facets of natural singing behaviours that account for reward-related variability in the singing experience, thereby helping to explain why some individuals enjoy singing more than others. We conducted three studies using factorial and correlational analyses to identify these dimensions of reward natural singing. In **Study 1**, we sought to determine, for the first time, which facets could explain variability in singing reward and enjoyment regardless of musical training or ability. Based on prior research on individual differences in music reward and engagement, we created a pool of items representing various dimensions of the natural singing experience, which were administered to a large international sample of English proficient participants (n = 606). Exploratory factor analysis (EFA) was conducted to identify the latent facets explaining variability in singing reward. As a result, we developed the Barcelona-Aarhus Natural Singing Engagement Questionnaire (BANSEQ) that assess sensitivity to singing reward across five distinct facets. In **Study 2**, the BANSEQ was administered to a second large international sample (n = 430) and confirmatory factor analysis (CFA) was used to validate the reliability and factor structure of singing reward. Finally, in **Study 3**, we examined the shared variance between the identified singing reward facets from the BANSEQ and previously validated constructs of music reward and engagement, and we explored sociodemographic and other psychological factors that are associated to each facet.

## Methods

### Study 1. Facets underlying sensitivity to reward and enjoyment in natural singing

#### Aim and procedures

The aim of the first study was to identify the facets that explain variability in sensitivity to deriving reward and enjoyment from natural singing in the general population. We initially created a pool of 84 items covering various aspects of natural singing, defined here as non-professional singing behaviours occurring in daily life. Accordingly, the items were designed to be independent of individuals’ prior singing training and their level of singing or musical skills. Item content was informed by statements drawn from online forums for amateurs singers, and from three existing questionnaires assessing musical reward sensitivity and musical engagement: the extended version of the Barcelona Music Reward Questionnaire (eBMRQ; Cardona et al., 2022; Mas-Herrero et al., 2013), the Music Engagement Questionnaire (MusEQ; Vanstone et al., 2016) and the Goldsmith Musical Sophistication Index (Gold-MSI; Müllensiefen et al., 2013). The 84 items were initially categorized into seven domains based on the subscales of the aforementioned questionnaires: *pleasure and emotion evocation* (i.e., the degree of pleasant or emotional responses to singing), *mood regulation* (i.e., changes in mood when singing or the habit of singing to change mood), *social reward* (i.e., enjoyment when singing in a group), *explorative component* (i.e., the desire to learn new songs or acquire material to sing or record), *absorption* (i.e., feeling uplifted when singing alone or in a group, or feeling connected with singers or their lyrics), *singing frequency* (i.e., the amount of time spent singing in daily life), and *singing responsiveness* (i.e., the desire to start singing or join group singing). In addition, six of the 84 items were inverted to control for acquiescence bias. Each statement was formulated so that respondents could indicate the degree to which they identified with it, using a 7-point Likert scale: (1) “*strongly disagree*”, (2) “*disagree*”, (3) “*somewhat disagree*”, (4) “*neither agree nor disagree*”, (5) “*somewhat agree*”, (6) “*agree*” and (7) “*strongly agree*”.

An online survey including the singing-related items was administered via the Qualtrics platform to an international sample of adult participants with advanced or proficient English. In addition, participants provided sociodemographic information and details of their prior musical and singing experience, and they completed two questionnaires assessing musical hedonism and musical engagement (for further details, see the “Instruments” section of Study 3). All participants provided informed consent prior to beginning the survey.

#### Participants

A total of 606 adult volunteers (73% women, mean age = 38.1 ± 13.3 years old) responded to the survey in English. All participants reported advanced or proficient written and oral English skills. Of these participants, 77.3% reported having prior musical experience (mean years of training = 10.3 ± 9.4). The international sample represented 82 different nationalities (see *Supplementary Table 1*) and ten nationality categories based on the Global Clustering of countries by Culture (Mensah & Chen, 2012): Latin European (39.3%), Anglo-Saxon (27.3%), Latin American (9.1%), Nordic (9.1%), German (5.3%), Eastern European (3.5%), South-East Asian (2.5%), Confucian (1.3%), Middle Eastern (1.3%), and African (1%).

#### Data analysis

##### Sample and correlation matrix assessment

To evaluate whether the sample size was appropriate for factor analysis, we applied the Seneca Estimate method (Lorenzo-Seva & Ferrando, 2024). This method estimates the required sample size to reproduce the expected population correlation matrix within a specified precision threshold. We set a threshold of .003 for the Root Mean Square of Residuals (RMSR) between (1) the observed correlation matrix in the sample and (2) the population correlation matrix. If the actual sample size is close to the value suggested by the Seneca Estimate, it can be considered adequate for the desired level of precision.

##### Initial selection of items

Exploratory factor analysis (EFA) is widely employed for item analysis during the initial stages of scale development, particularly when working with large item pools, as in our case (84 items). Under such conditions, the inclusion of inappropriate or ineffective items can hinder the analysis, making it more difficult to determine the underlying dimensionality and structure of the scale.

To address this issue and identify the most suitable items prior to specifying any factor model, we applied Gulliksen’s pool technique. This straightforward procedure is designed to detect potentially problematic items before committing to a particular factorial solution (Ferrando et al., 2023). Specifically, the method defines regions of item appropriateness and efficiency based on the influence of two key item characteristics: extremeness and consistency.

##### Exploratory Factor Analysis

An exploratory factor analysis was performed to determine the optimal number of factors and explore whether the proposed facets of the test were adequately represented by a single factor. To assess the adequacy of matrix correlation for factor analysis, the Kaiser-Meyer-Olkin (KMO) Test of Sampling Adequacy was computed (Kaiser & Rice, 1974). In addition, item-level normed measures of sampling adequacy (Normed-MSA) were inspected to identify items with sufficient shared variance. Normed-MSA values below .50 indicate that an item does not adequately measure the same construct as the remaining items and should therefore be removed (Lorenzo-Seva & Ferrando, 2021).

To evaluate dimensionality, we conducted parallel analysis (PA; Timmerman & Lorenzo-Seva, 2011) and computed essential unidimensionality indices (Ferrando & Lorenzo-Seva, 2018): unidimensional congruence (UniCo), explained common variance (ECV), and the mean of item residual absolute loadings (MIREAL). PA compares the factor structure of the observed data with that of randomly generated datasets under a null model. The number of factors recommended for extraction corresponds to the number of dimensions in the observed data that account for more common variance than those in the random datasets. The essential unidimensionality indices were used to assess whether a strong, dominant factor is present. A unidimensional solution is supported when all three indices meet their respective thresholds: UniCo > 0.95, ECV > 0.85, and MIREAL < 0.30.

We expected a factor model in which each item was associated with one of the five proposed facets, while the five facets were sufficiently correlated to justify an overall individual score. The factor analysis solution was estimated using the Unweighted Least Squares (ULS) criterion, and the chi-square statistic was scaled with LOSEFER (Lorenzo-Seva & Ferrando, 2023). While the standard approach scales the chi-square statistic based on the mean and variance of its distribution, LOSEFER further incorporates skewness and kurtosis, leading to a more accurate scaled chi-square value. To assess the goodness of fit of the factor model, we examined the following indices: Root Mean Square Error of Approximation (RMSEA; values between .050 and .080 indicate a fair fit), Comparative Fit Index (CFI; values above .98 indicate excellent fit), Goodness of Fit Index (GFI; values above .98 are recommended for a good fit), and Root Mean Square of Residuals (RMSR; with a dataset-adaptive threshold proposed by Kelly) (Harman, 1976). Although we expected the pool of items to be essentially unidimensional, we also extracted the number of factors suggested by the number of facets that we expected and applied Promin rotation (Lorenzo-Seva, 1999) to assess whether the facets were clearly defined.

Factor analyses were conducted using FACTOR software (Lorenzo-Seva & Ferrando, 2006), and descriptive analyses were performed using RStudio (version 2024.04.2). Statistical significance was set at a two-sided *p <* .005.

### Study 2: Confirmatory factorial analysis of the Barcelona-Aarhus Natural Singing Engagement Questionnaire (BANSEQ)

#### Aim and procedures

The aim of the second study was to conduct a confirmatory factor analysis (CFA) of the psychometric tool obtained in Study 1 to measure sensitivity to natural singing reward, which we called Barcelona-Aarhus Natural Singing Engagement Questionnaire (BANSEQ; see *Appendix A*). As the outcome of the first exploratory study suggested a unidimensional higher-order factor structure comprising 20 items across five dimensions, we assessed this factor model using CFA. The five facets are: (1) *pleasure and emotional evocation* (i.e., the degree of pleasure or emotional response to singing), (2) *social singing reward* (i.e., the enjoyment derived from singing with others), (3) *mood regulation through singing* (i.e., the use of singing to manage emotions or improve mood), (4) *singing frequency* (i.e., how often one engages in everyday singing), and (5) *inattentional singing during routine tasks* (i.e., the tendency to sing automatically or without explicit awareness during routine tasks).

We administered the BANSEQ in an online survey to a new international sample of adult participants with advanced or proficient English. The survey also included questions on sociodemographic information and prior musical and singing experience, as well as six questionnaires assessing reward-related responses to musical, social, and sensory experiences, musical engagement, self-perceived singing abilities, responsiveness to aesthetic experiences, and personality traits (for further details, see the “Instruments” section of Study 3).

The survey was administered via the Qualtrics platform and distributed across English-speaking countries or countries with high levels of English proficiency. All eligible individuals with proficient English were invited to participate voluntarily, with the option of entering a draw for three electronic tablets once the survey closed. Before beginning the survey, participants provided informed consent.

#### Participants

A total 430 adult volunteers (60% women, mean age = 38.1 ± 13.6 years) completed the online survey in English. All participants reported advanced or proficient written and oral English skills (70.60% native English speakers, 21.30% English as a second language, 6.71% English as a third language). Of these, 62.2% reported prior musical experience (mean years of training = 7.8 ± 7.7) and represented the following nationality categories based on the Global Clustering of Countries by Culture (Mensah & Chen, 2012): Anglo-Saxon (68.1%), Latin European (13%), Latin American (4.2%), Eastern European (3.7%), Nordic (2.6%), South-East Asian (2.6%), German (1.6%), African (1.2%), Confucian (1.2%), and Middle Eastern (1.2%).

To evaluate whether the sample size was appropriate for factor analysis, we applied the Seneca Estimate method (Lorenzo-Seva & Ferrando, 2024). This method estimates the required sample size to reproduce the expected population correlation matrix within a specified precision threshold. We set a threshold of .003 for the Root Mean Square of Residuals (RMSR) between (1) the observed correlation matrix of the sample and (2) the population correlation matrix. If the actual sample size is close to the value suggested by the Seneca Estimate, it can be considered adequate for the desired level of precision.

#### Data analysis

##### Confirmatory Factor Analysis

A confirmatory factor analysis (CFA) was conducted to evaluate whether the unidimensional factor model observed in Study 1 would generalize to the population level. We conducted a Mild-Restricted CFA (MRCFA; Lorenzo-Seva, 2025), since traditional CFA is increasingly regarded as a less appropriate (Steenkamp & Maydeu-Olivares, 2023) due to its tendency to converge on biased factor parameters. MRCFA mitigates this issue by first estimating the free parameters of the model before imposing the model restrictions, which reduces the likelihood of bias in the estimated parameters. The extraction method used was ULS, and the chi-square statistic was scaled using LOSEFER. To assess the goodness of fit of the factor model, we examined the following indices: RMSEA, CFI, GFI, and RMSR.

##### Factor Analysis of the total sample

A final factor analysis was performed by combining the samples of Studies 1 and 2 (n = 1036) in order to obtain the most accurate estimates of item loadings in the factor model and to compute factor scores. A factor analysis solution was fitted by ULS to retain a single factor. To estimate factor reliabilities of factor scores, we computed ORION reliability index of factor scores: values above .80 are considered acceptable. The quality of the factor solution was inspected using construct replicability, and the quality of factor scores estimates were assessed with indices H, and Sensitivity Ratio (SR-index) (Ferrando & Lorenzo-Seva, 2018). The H index evaluates how well a set of items represents a common factor. It is bounded between 0 and 1 and approaches unity as the magnitude of the factor loadings and/or the number of items increases. High H values (>.80) suggest a well-defined latent variable. The SR-index can be interpreted as the number of different factor levels that can be differentiated on the basis of the factor score estimates. If factor scores are to be used for individual assessment, marginal reliabilities above .80, and SR above 2 are recommended.

Factor analyses were conducted using FACTOR software (Lorenzo-Seva & Ferrando, 2006), and descriptive analyses were performed using RStudio (version 2024.04.2). Statistical significance was set at a two-sided *p*-value < .005.

### Study 3: Convergent validity of BANSEQ and associated factors

#### Aim and procedures

The aim of the third study was to examine the convergent validity of the BANSEQ using previously validated measures of other musical behaviours and hedonic capacity toward musical, social, and sensory stimuli and experiences, as well as to assess the influence of sociodemographic and psychological characteristics in modulating sensitivity to natural singing reward across its distinct facets. To this end, we combined the samples from Studies 1 and 2, compared singing reward-related responses across sociodemographic variables, and correlated them with the remaining continuous measures.

#### Participants

A total of 1036 participants (from Studies 1 and 2) completed the BANSEQ (67.5% women, mean age = 38.1 ± 13.4 years). Of this sample, 71.1% reported prior musical experience (mean years of training = 9.4 ± 8.9) and represented ten nationality categories based on the Global Clustering of Countries by Culture (Mensah & Chen, 2012): Anglo-Saxon (68.1%), Latin European (13%), Latin American (4.2%), Eastern European (3.7%), Nordic (2.6%), South-East Asian (2.6%), German (1.6%), African (1.2%), Confucian (1.2%), and Middle Eastern (1.2%).

#### Instruments

In addition to the BANSEQ, participants completed the following self-report scales:

1. The <u>eBMRQ</u> was included in both Studies 1 and 2 to evaluate participants’ capacity to derive reward from music-related activities. This questionnaire includes 24 items classified into six facets (*musical seeking*, *emotion evocation*, *mood regulation*, *social reward*, *sensory-motor*, and *absorption*). The reliability coefficients for the six facets range from 0.84 to 0.93, and the reliability for the overall scale is 0.95 (Cardona et al., 2022; Mas-Herrero et al., 2013).
2. The <u>Snaith-Hamilton Pleasure Scale</u> (SHAPS; Snaith et al., 1995) was included in Study 2 to measure general hedonic capacity across four domains of pleasurable responses beyond music (*interest / pastimes*, *social interaction*, *sensory experiences*, and *food / drink*). The scale includes 14 items and has a reliability of 0.86.
3. The <u>Music Engagement Questionnaire</u> (MusEQ; Vanstone et al., 2016) was included in Study 2 to assess participants’ engagement with music in daily life. It contains 35 items across six facets (*daily*, *emotion*, *performativity*, *responsivity*, *preference,* and *consumption*). Subscale reliabilities range from 0.65 to 0.82, and the reliability for the overall scale is 0.92.
4. The <u>Gold-MSI Singing Ability subscale</u> (Müllensiefen et al., 2013) along with the behavioural subscale of the <u>Seattle Singing Accuracy Protocol</u> (SSAP; Demorest & Pfordresher, 2015; Scholtz et al., 2021) were included in both Studies 1 and 2 to evaluate participants’ self-perceived singing ability. The Gold-MSI Singing Ability subscale consists of 7 items and has a reliability of 0.87. The SSAP behavioural subscale consists of three items measuring both perceived skills and external feedback on singing, and the full protocol shows a reliability of 0.92. Both subscales were combined to obtain a total score of Perceived Singing Abilities (PSA).
5. The <u>Aesthetic Responsiveness Assessment</u> (AReA; Scholtz et al., 2021) was included in Study 2 to measure responsiveness to aesthetic experiences beyond music (e.g., visual arts, dance and performance). This recently developed questionnaire contains 14 items grouped into three facets (*aesthetic appreciation*, *intense aesthetic experience*, and *creative behaviour*). Reliability coefficients for the facets are 0.84, 0.80, and 0.63, respectively, with an overall reliability of 0.82.
6. The extra-short form of the <u>Big Five Inventory-2</u> (BFI-2-XS; Soto & John, 2017) was included in Study 2 to assess participants’ personality traits across 15 items classified into five dimensions (*extraversion*, *agreeableness*, *conscientiousness*, *neuroticism*, and *openness*), with reliabilities ranging from 0.51 to 0.72.

#### Data analysis

##### Convergent validity of BANSEQ

Correlation analyses were conducted between the BANSEQ total and facet scores and the eBMRQ (total and facet scores), SHAPS (total score) and MusEQ (total and facet scores). We conducted Spearman rank-order correlations because the BANSEQ total and facet scores showed non-normal distribution. Bonferroni corrections for multiple comparisons were applied when comparing several subscales simultaneously and for post-hoc comparisons.

##### Factors associated with BANSEQ

The effects of sociodemographic variables, prior musical expertise, self-perceived singing ability, responsiveness to aesthetic stimuli, and personality traits on natural singing engagement were examined using correlation and descriptive analyses. Correlations were computed between the BANSEQ total and subscale scores and age, years of musical training, PSA (total score), AReA (total and facet scores), and BFI-2-XS (dimension scores). Spearman correlations were applied due to the non-normal distribution of BANSEQ scores. A Kruskal-Walli’s test was applied to assess gender differences (women, men, and non-binary participants) in natural singing engagement (BANSEQ total and facet scores)

All descriptive and correlational analyses were conducted using RStudio (version 2024.04.2). Statistical significance was evaluated using a two-sided *p* < .005.

## Results

### Study 1: Facets underlying sensitivity to reward and enjoyment in natural singing

#### Sample and correlation matrix assessment

The Seneca Estimate suggested that a sample size of 610 participants was required, with our sample (n = 606) closely meeting this requirement. The inter-item correlation matrix for this sample yielded a Kaiser-Meyer-Olkin (KMO) value of .922 (bootstrap 95% confidence interval: .895–.9925), with normed measures of sampling adequacy (MSA) values for individual items ranging from .893 to .938. These results indicate that the correlation matrix was suitable for factor analysis and the 20 items contributed substantially to the shared variance.

#### Initial selection of items

To ensure that the five facets were equally represented in the final item set, we applied Gulliksen’s pool technique separately to each of the seven domains previously used to classify the full pool of 84 items (i.e., the analysis was conducted seven times). Inspection of the results obtained from Gulliksen’s pool technique enabled us to identify subsets of items with extremeness values ranging between .10 and .90, and consistency values between .20 and .75.

In two of the domains, none of the items met the criteria for inclusion in a factor analysis. In contrast, within the remaining five domains (Singing Frequency; Social Reward; Pleasure and Emotional Evocation; Mood Regulation; and Inattentional Singing), we identified four items per domain that satisfied the adequacy criteria for factor analysis. This yielded a final set of 20 items, which were subsequently included in the exploratory factor analysis. Of the initial seven domains considered in the development of the 84-item pool, only five were ultimately considered to define the hedonic experience of singing.

#### Exploratory Factor Analysis

Parallel analysis (PA) suggested that a single-factor model comprising 20 items should be considered for the sample data. When a single factor was extracted, the goodness-of-fit indices did not indicate a perfect fit but fell within an acceptable range: RMSEA = .081 (bootstrap 95% confidence interval: .080–.083), CFI = .955 (bootstrap 95% confidence interval: .951–.958), GFI = .954 (bootstrap 95% confidence interval: .932–.962), and RMSR = .093 (bootstrap 95% confidence interval: .084–.098). Additionally, the essential unidimensionality indices supported the presence of a dominant factor: UniCo = .931 (bootstrap 95% confidence interval: .909–.961), ECV = .830 (bootstrap 95% confidence interval: .808–.859), and MIREAL = .223 (bootstrap 95% confidence interval: .194–.243).

To assess whether the five domains were actually contributing to defining the hedonic experience of singing, we also explored a five-factor model. The extracted five-factor solution demonstrated an excellent fit, with goodness-of-fit indices as follows: RMSEA = .025, CFI = .997, GFI = .998, RMSR = .0162, and Kelley’s criterion = .0407. Therefore, this unidimensional solution, which measures overall sensitivity to singing reward, comprises five distinct factors that explain the variability in singing reward. Based on these results, we developed the Barcelona-Aarhus Natural Singing Engagement Questionnaire (BANSEQ), a psychometric tool capturing these five facets. Rotated factor loadings supported the adequate representation of each facet within the model (see Supplementary Table 2). Correlations between factors ranged from .346 to .630 (see Supplementary Table 3), demonstrating the model’s essential unidimensionality. The BANSEQ comprises 20 items classified into five facets (with four items per facet):

- The *pleasure and emotional evocation* facet assess the degree of consummatory pleasure or emotional response experienced when singing individually, capturing natural singing behaviours that occur in solitary contexts (e.g., singing in the shower or in the car). It includes items such as “*I feel happy when singing by myself*” and “*I learn to sing songs that I love regardless of whether I will show them to someone else or not*.”
- The *mood regulation through singing* facet reflects the habitual use of singing to influence or manage one’s emotional state, whether individually or in a group, and includes items such as “*When I am sad, I sing certain songs to feel better*” and “*When someone hurts my feelings, I sing songs to deal with it*”.
- The *social singing reward* facet addresses the inherently social nature of singing and measures the extent to which individuals enjoy singing in social contexts. It includes items such as “*If others are singing, I join in*,” as well as reverse-scored items like “*I hate social activities related to singing (e.g., going to karaoke)*”.
- The *singing frequency* facet quantifies the level of non-professional singing engagement by assessing how often individuals sing voluntarily in their daily lives, with items such as “*I sing weekly alone or in a group”* and “I have practiced singing on a regular basis during my lifetime”.
- The *inattentional singing during routine tasks* facet captures the tendency to engage in automatic or habitual singing without explicit awareness while performing chores, routine activities, or movement-related tasks, functioning as a form of intrinsic entertainment.

Example items include “I sing or hum as I go about my daily activities” and “I sing or hum when doing certain tasks (e.g., cooking or cleaning)”.

### Study 2: Confirmatory factorial analysis of the Barcelona-Aarhus Natural Singing Engagement Questionnaire (BANSEQ)

#### Sample and correlation matrix assessment

The Seneca Estimate applied to a confirmatory model with one expected factor indicated that a sample of 340 participants would be required, with our sample (n = 430) exceeding this requirement. The inter-item correlation matrix for the second sample yielded a KMO value of .947 (bootstrap 95% confidence interval: .895–.9925), indicating that this correlation matrix was suitable for factor analysis and that all items contributed substantially to the shared variance.

#### Confirmatory Factor Analysis of the second sample

The factor model assessed in the CFA was the unidimensional model. The goodness-of-fit indices indicated that the model fit was within an acceptable range: RMSEA = .063 (bootstrap 95% confidence interval: .056 - .064), CFI = .985 (bootstrap 95% confidence interval: .984 - .988), GFI = .938 (bootstrap 95% confidence interval: .935 - .949), and RMSR = .080 (bootstrap 95% confidence interval: .070 - .085). All facets showed satisfactory loadings on the factor (see *Supplementary Table 4*), and inter-factor correlations ranged from .488 to .841 (see *Supplementary Table 5*), further supporting the model’s essential unidimensionality. These results suggest that the unidimensional factor model can be expected to generalise to the population level.

#### Factor Analysis of the total sample

Since the analyses of samples from Studies 1 and 2 led to the same conclusions, the samples were combined (n = 1036) and jointly analysed to assess the quality of the factor solution and to compute factor scores for each participant. As shown in *Table 1*, factor loadings indicated adequate representation of all facets in the model. Additionally, the inter-factor correlation matrix revealed moderate-to-strong correlations ranging from .442 to .747 (see *Table 2*), confirming the appropriateness of the unidimensional model. The ORION reliability index was .941, indicating that the factor scores had an acceptable level of reliability. Finally, the indices used to evaluate the quality of the factor solution suggested a well-defined latent variable (see *Table 3*).

**Table 1.** Factor loadings for the unidimensional solution and item loadings across facets for the whole sample in Study 2 (n = 1036). Salient loadings are printed in bold. (BANSEQ: Barcelona-Aarhus Natural Singing Engagement Questionnaire; SF: Singing Frequency; SR: Social Reward; PE: Pleasure and Emotional evocation; MR: Mood Regulation; IS: Inattentional Singing).

| Item |  | BANSEQ | SF | SR | PE | MR | IS |
| --- | --- | --- | --- | --- | --- | --- | --- |
| 1 | *I sing weekly alone or in a group.* | **.648** | **.556** | .108 | .224 | -.166 | .081 |
| 2 | *I become cheerful when I sing in a group.* | **.610** | .031 | **.749** | .125 | -.028 | -.014 |
| 3 | *I enjoy singing.* | **.780** | .202 | .237 | **.715** | -.101 | -.090 |
| 4 | *When I am sad, I sing certain songs to feel better.* | **.651** | .016 | -.020 | -.004 | **.892** | -.032 |
| 5 | *I sing or hum as I go about my daily activities.* | **.691** | .033 | .037 | -.109 | .000 | **.934** |
| 6 | *I have practiced singing on a regular basis during my lifetime.* | **.624** | **.881** | .046 | -.037 | -.046 | -.034 |
| 7 | *If others are singing, I join in.* | **.618** | -.017 | **.685** | -.059 | .113 | .150 |
| 8 | *I sing by myself for pleasure.* | **.806** | -.005 | -.109 | **.816** | .070 | .126 |
| 9 | *Singing takes me back to my teen years.* | **.498** | -.090 | .075 | .058 | **.477** | .117 |
| 10 | *I sing or hum when doing certain tasks (e.g., cooking or cleaning).* | **.668** | -.111 | -.022 | .227 | -.144 | **.854** |
| 11 | *There are times when I do not nothing except singing.* | **.636** | **.631** | -.053 | -.031 | .112 | .167 |
| 12 | *I feel uncomfortable when signing in a group.* | **-.429** | -.110 | **-.817** | .105 | .110 | .040 |
| 13 | *I feel happy when singing by myself.* | **.748** | -.146 | -.094 | **.999** | .021 | .026 |
| 14 | *When someone hurts my feelings, I sing songs to deal with it.* | **.619** | .129 | -.016 | .012 | **.769** | -.055 |
| 15 | *I sing or hum while hiking or travelling.* | **.648** | .052 | .062 | .029 | .003 | **.663** |
| 16 | *I do sing or practice singing for a certain amount of time per day.* | **.653** | **.785** | -.055 | .136 | .003 | -.057 |
| 17 | *I hate social activities related to singing (e.g., going to karaoke).* | **-.414** | .179 | **-.631** | -.068 | -.116 | .023 |
| 18 | *I learn to sing songs that I love regardless of whether I will show it to someone else or not.* | **.716** | .235 | -.048 | **.581** | .046 | .016 |
| 19 | *I become emotional when I sing certain songs.* | **.660** | -.155 | .092 | .552 | **.385** | -.111 |
| 20 | *If I am bored, I sing or hum a song to pass the time (e.g., performing chores, waiting in a queue, etc.)* | **.659** | .068 | -.013 | -.117 | .076 | **.833** |

**Table 2.** Inter-factor correlation matrix for the whole sample in Study 2 (n = 1036). (SF: Singing Frequency; SR: Social Reward; PE: Pleasure and Emotional evocation; MR: Mood Regulation; IS: Inattentional Singing).

|  | SF | SR | PE | MR |
| --- | --- | --- | --- | --- |
| **SR** | .598 |  |  |  |
| **PE** | .747 | .632 |  |  |
| **MR** | .635 | .442 | .696 |  |
| **IS** | .632 | .443 | .717 | .651 |

**Table 3.** Indices to assess the quality of the factor solution related to the total sample.

| Index | BANSEQ |
| --- | --- |
| H | .940 |
| SR | 3.978 |
| EPTD | 94.7% |

To analyse participants’ scores on the factor, factor scores should be computed rather than simply summing item responses. Computing factor scores provides a more accurate representation of the underlying traits or abilities measured by the test, as it accounts for the relationships between items and the latent variables they intend to capture. The statistics of factors scores of the total sample are provided in *Table 4*. In the supplementary materials, we provide researchers with an Excel file that calculates factor scores from participants’ item responses (see *Supplementary Material*). Additionally, the raw data are freely available on the Open Science Framework (OSF).

**Table 4.** Descriptive statistics of BANSEQ factor scores for the combined samples 1 and 2, presented for each factor and for the overall BANSEQ score, as well as the raw additive BANSEQ mean score with standard deviation (SD).

| BANSEQ facets | 1<sup>st</sup> and 2<sup>nd</sup> sample<br>(n = 1036) |  |
| --- | --- | --- |
|  | Mean | SD |
| Pleasure and emotional evocation | 50.00 | 10.01 |
| Social reward | 50.00 | 10.00 |
| Singing frequency | 49.99 | 10.02 |
| Mood regulation | 49.99 | 10.02 |
| Inattentional singing | 50.01 | 10.02 |
| Overall Singing Reward (BANSEQ) | 50.00 | 10.00 |
| Overall Singing Reward ($\Sigma$ items) | 91.49 | 20.65 |

### Study 3: Convergent validity of BANSEQ and associated factors

#### Convergence validity with other dimensions of reward sensitivity

Correlations of the BANSEQ (total and facet scores) with the eBMRQ (total and facet scores) and the SHAPS (total score) are presented in *Table 5*. All BANSEQ facets and the total score correlated positively with all eBMRQ facets and total scores. The strongest association for the overall BANSEQ score was with the overall eBMRQ score, sharing nearly 34% of the variance (*r* = .58; *p* < .001). Among the BANSEQ facets, *mood regulation through singing* showed the strongest associations with the total eBMRQ score (r = .53; *p* < .001), and the second strongest association within this facet was observed with *mood regulation* through music listening (*r* = .46; *p* < .001). Regarding *social singing reward*, the eBMRQ facet most strongly associated was also *social reward* (*r* = .42; *p* < .001). With respect to general hedonic capacity, levels of anhedonia measured with the SHAPS correlated negatively with all BANSEQ facets and total score, and the strongest negative correlation was observed for the BANSEQ *inattentional singing* facet (*r* = - .34; *p* < .001), followed by the BANSEQ total score (*r* = -.31; *p* < .001). In contrast, the weakest association with the SHAPS was found for the BANSEQ *social singing reward* facet (*r* = -.17; *p* < .001).

**Table 5.**
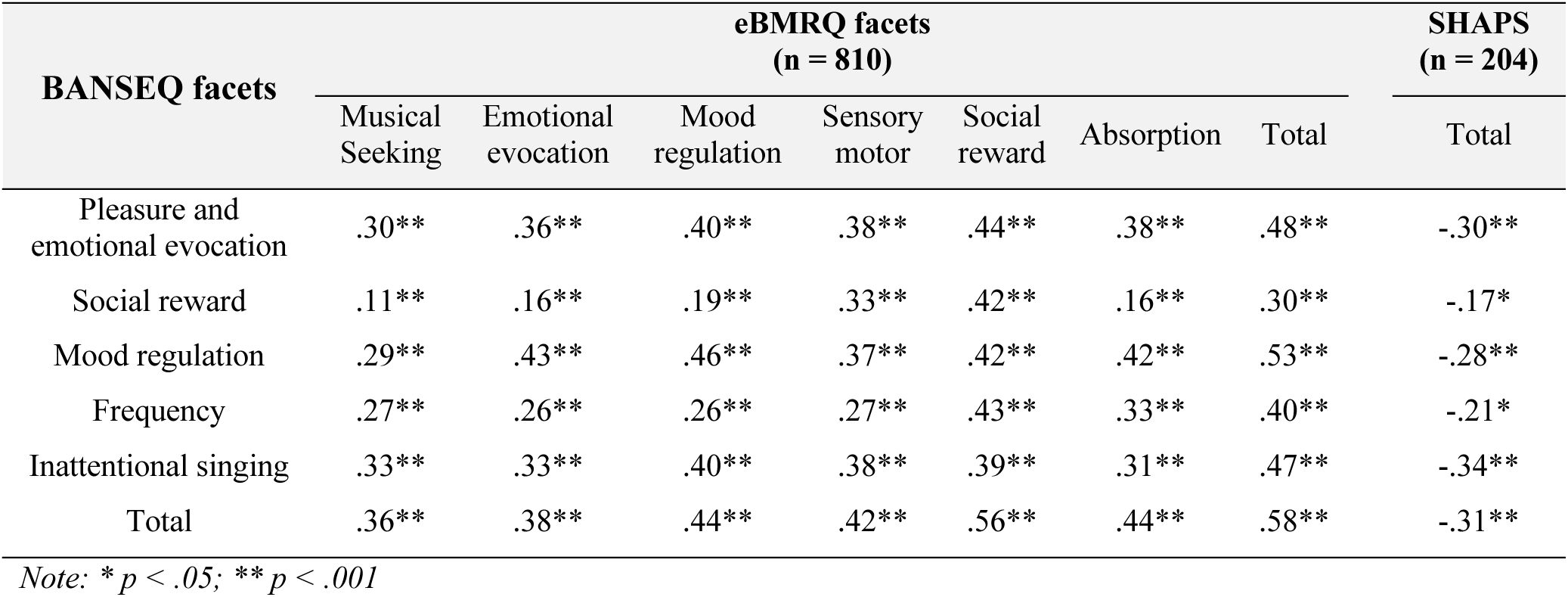
Correlations between the BANSEQ total and facet scores with the eBMRQ and the SHAPS scores. Spearman’s Rho (*r*) are presented per each comparison. (BANSEQ: Barcelona-Aarhus Natural Singing Engagement Questionnaire; eBMRQ: extended version of the Barcelona Music Reward Questionnaire; SHAPS: Snaith-Hamilton Pleasure Scale).

Since the BANSEQ and eBMRQ score distributions did not fully converge (shared variance = 34%), we plotted the scores to visually inspect the cross-distribution of both scales. First, overall scores on each scale were normalized using z-scores, setting the mean of each distribution to zero. These standardized scores were then plotted as x-y coordinates (x = BANSEQ; y = eBMRQ) in a heat map, where colour intensity reflects the number of participants in each cell (darker colours indicate a higher number of participants, and lighter colours indicate fewer participants). As shown in *Figure 1*, most participants (77.55%) fell within the normal range on both BANSEQ and eBMRQ, with overall scores between ±1.5 SD from the mean on each scale. However, 3.56% of individuals (marked by grey squares) reported low music enjoyment (eBMRQ overall score below -1.5 SD) while showing typical levels of singing enjoyment (BANSEQ overall score between ±1.5 SD). Conversely, 2.92% of participant exhibited the opposite pattern, reporting low singing enjoyment (BANSEQ overall score below -1.5 SD) while showing normal sensitivity to music reward (eBMRQ overall score between ±1.5 SD).

**Figure 1.**
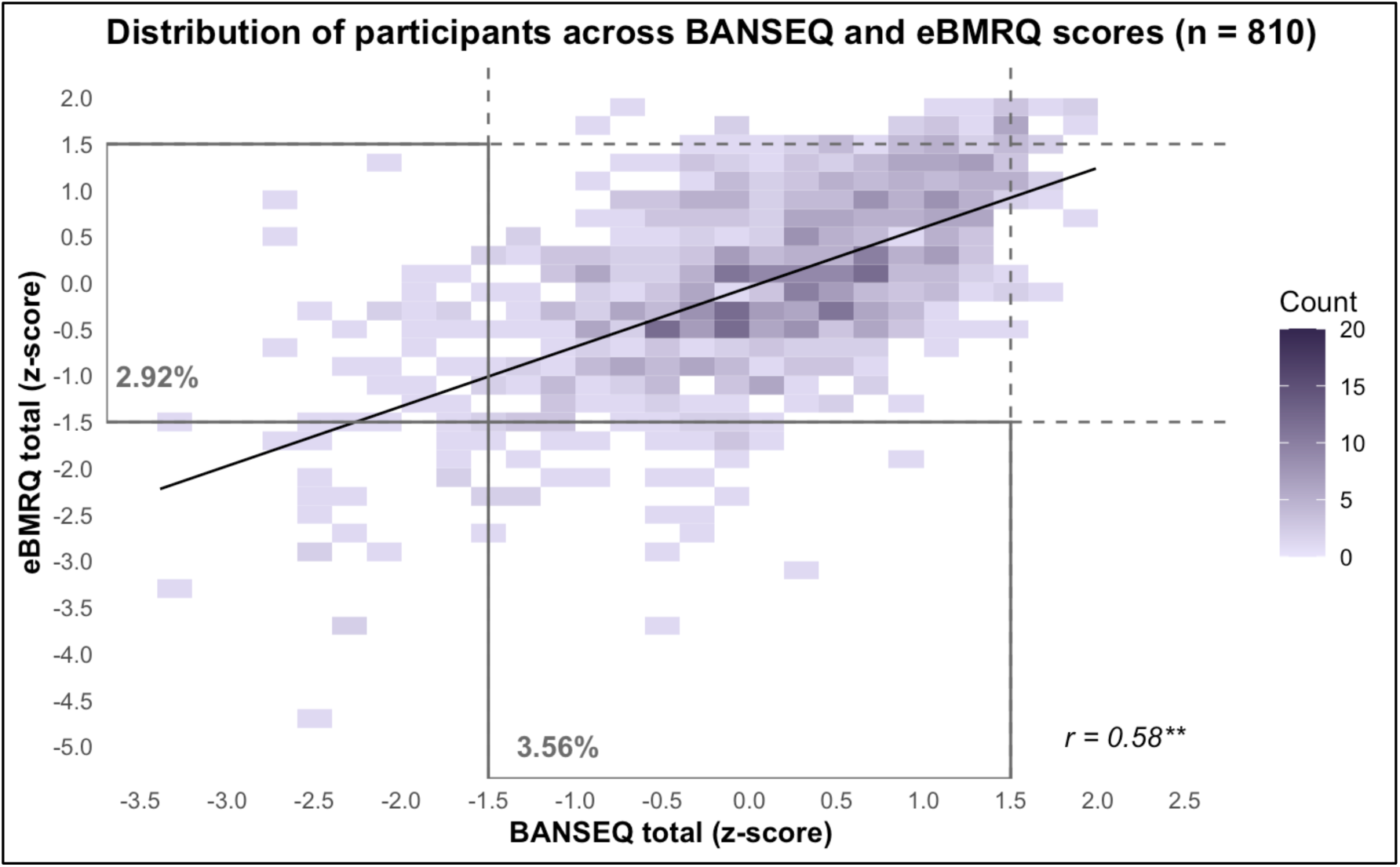
Distribution of BANSEQ and eBMRQ total scores in the third sample (Study 3). Scores were z-transformed and plotted as x-y coordinates in a heat map. Colour intensity reflects the number of participants per cell, with darker shades indicating higher frequencies. Dotted grey lines indicate cutoffs for values outside the normal range (±1.5 SD from the mean). Grey squares represent participants with low scores on one scale (below -1.5 SD) and scores within the normal range on the other scale (±1.5 SD from the mean). (BANSEQ: Barcelona-Aarhus Natural Singing Engagement Questionnaire; eBMRQ: extended version of the Barcelona Music Reward Questionnaire).

To further explore the dimensions of singing reward in participants showing a greater dissociation between sensitivity to general music reward and natural singing reward, we compared BANSEQ facet scores across different groups. Participants were classified as either those who preferred singing over general music engagement (P-Sing; characterized by normal BANSEQ scores and lower eBMRQ scores), and those who showed normal levels of general music enjoyment but did not particularly enjoy singing (P-Music; characterized by normal eBMRQ scores and lower BANSEQ scores). In addition, we classified the sample according to overall levels of singing enjoyment. Participants with particularly high levels of singing enjoyment (i.e., singing hyperhedonia) were labelled as High-Sing (overall BANSEQ scores above +1.5 SD from the mean), whereas those with particularly low levels of singing enjoyment (i.e., singing hypohedonia) were labelled as Low-Sing (overall BANSEQ scores below -1.5 SD from the mean). *Table 6* reports the mean BANSEQ overall scores and facet scores for each group (P-Sing, P-Music, High-Sing, and Low-Sing), together with the corresponding values for the whole sample. To facilitate visual comparison, the scores of the five BANSEQ facets were transformed onto a 1-5 scale. These values were plotted to compare the singing enjoyment profiles of the P-Sing and P-Music groups with that of the whole sample (see *Figure 2.A*), as well as to compare the profiles of the High-Sing and Low-Sing groups with the whole sample (see *Figure 2.B*).

**Figure 2.**
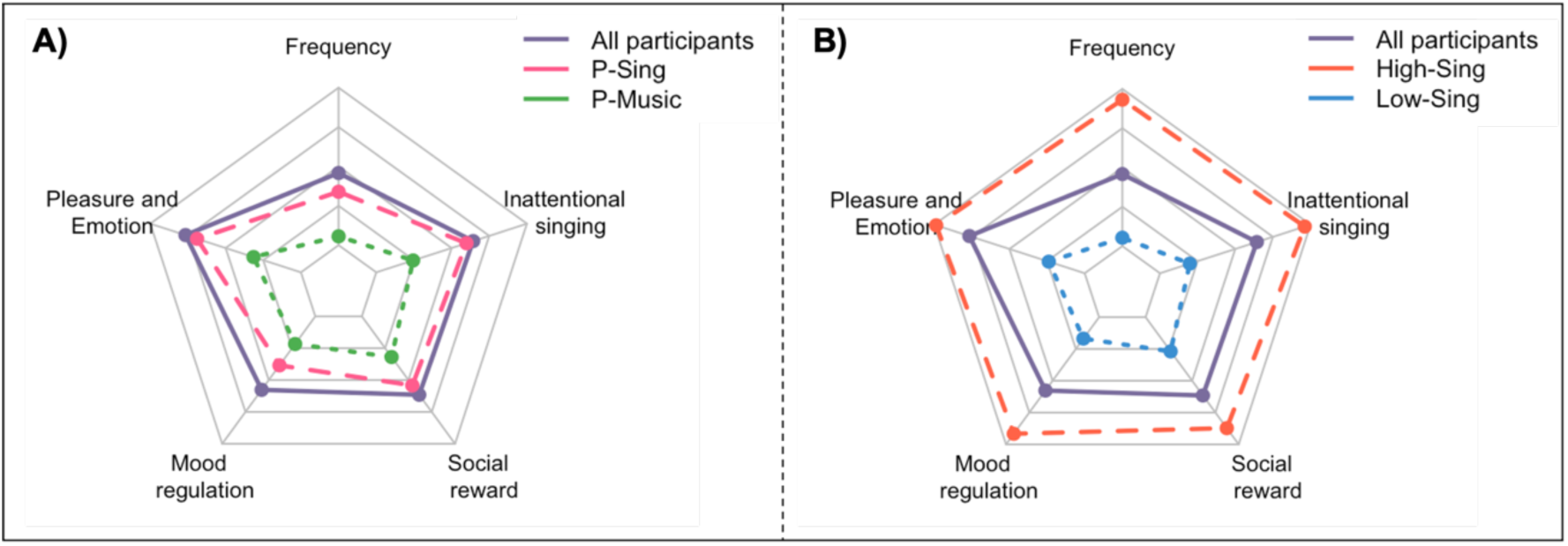
Graphical representation of BANSEQ profiles across participant groups. The star plots show the mean values of each BANSEQ facet, scaled to a 1-5 range, where 1 represents the minimum and 5 the maximum score observed in the sample. The profile of the whole sample (in purple) is compared with: (**A**) P-Sing (in pink) and P-Music participants (in green), and (**B)** High-Sing (in orange) and Low-Sing participants (in blue). (P-Sing: Preferring Singing, characterized by normal BANSEQ scores and lower eBMRQ scores; P-Music: Preferring Music, characterized by normal eBMRQ scores and lower BANSEQ scores; High-Sing: Higher Singing Enjoyment, characterized by higher BANSEQ scores; Low-Sing: Lower Singing Enjoyment, characterized by lower BANSEQ scores; BANSEQ: Barcelona-Aarhus Natural Singing Engagement Questionnaire; eBMRQ: extended version of the Barcelona Music Reward Questionnaire).

**Table 6.**
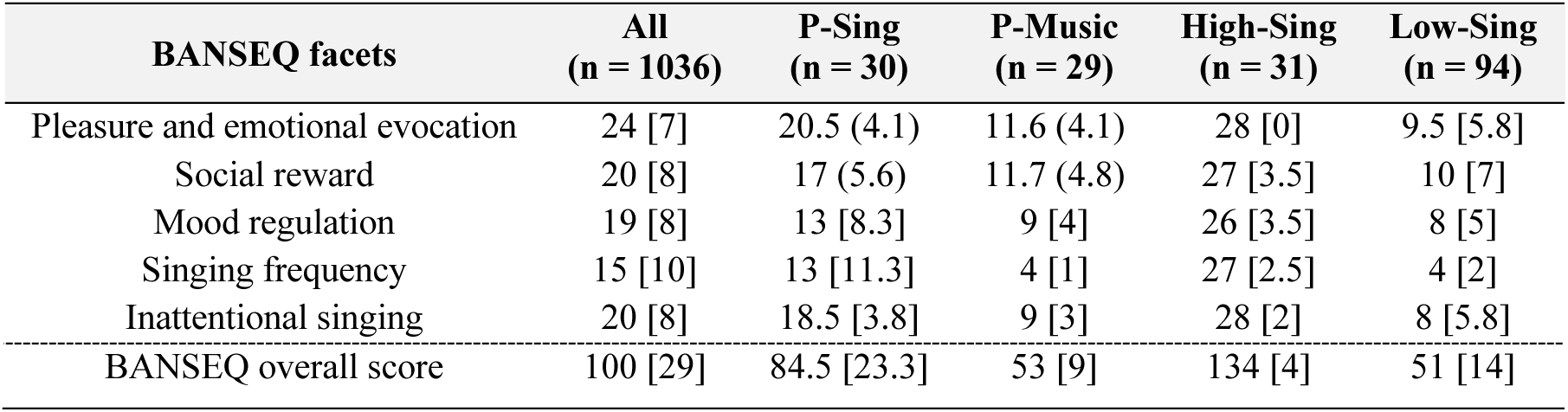
Descriptive statistics of BANSEQ across groups. Means with standard deviations (SD) or medians with interquartile ranges [IQR] of the BANSEQ overall score and each facet are reported for each participant group, as appropriate for parametric or nonparametric distributions. (All: All participants; High-Sing: Higher Singing Enjoyment, characterized by higher BANSEQ scores; Low-Sing: Lower Singing Enjoyment, characterized by lower BANSEQ scores; P-Sing: Preferring Singing, characterized by normal BANSEQ scores and lower eBMRQ scores; P-Music: Preferring Music, characterized by normal eBMRQ scores and lower BANSEQ scores; BANSEQ: Barcelona-Aarhus Natural Singing Engagement Questionnaire; eBMRQ: extended version of the Barcelona Music Reward Questionnaire).

| BANSEQ facets | All<br>(n = 1036) | P-Sing<br>(n = 30) | P-Music<br>(n = 29) | High-Sing<br>(n = 31) | Low-Sing<br>(n = 94) |
| --- | --- | --- | --- | --- | --- |
| Pleasure and emotional evocation | 24 [7] | 20.5 (4.1) | 11.6 (4.1) | 28 [0] | 9.5 [5.8] |
| Social reward | 20 [8] | 17 (5.6) | 11.7 (4.8) | 27 [3.5] | 10 [7] |
| Mood regulation | 19 [8] | 13 [8.3] | 9 [4] | 26 [3.5] | 8 [5] |
| Singing frequency | 15 [10] | 13 [11.3] | 4 [1] | 27 [2.5] | 4 [2] |
| Inattentional singing | 20 [8] | 18.5 [3.8] | 9 [3] | 28 [2] | 8 [5.8] |
| BANSEQ overall score | 100 [29] | 84.5 [23.3] | 53 [9] | 134 [4] | 51 [14] |

#### Convergence with general musical engagement

Correlations between the BANSEQ (total and facet scores) and the MusEQ (total and facet scores) are presented in *Table 7*. A moderate correlation was observed between the BANSEQ and MusEQ overall scores, indicating nearly 30% shared variance (*r* = .54; *p* < .001) (see *Figure 3.A*). The overall score of the BANSEQ correlated positively with all facets of the MusEQ, except for *preference* (i.e., related to how restricted or broad individuals’ music taste is), which only correlated weakly with the *social reward* BANSEQ facet (*r* = -.14; *p* < .037). The strongest association was between the BANSEQ total score and the MusEQ *responsivity* facet (*r* = .65; *p* < .001), which contains three items on inattentive, social, and individual singing or humming, and one item on body movement in response to music. Consistent with this, the strongest associations across BANSEQ facets were also observed with the MusEQ *responsivity* facet, with the BANSEQ *pleasure and emotional evocation* facet showing the strongest association (*r* = .60; *p* < .001). The *daily* facet of the MusEQ, which refers to everyday music listening, correlated positively with all BANSEQ facets except for *social singing reward*. The *emotion* facet of the MusEQ, which refers to emotional evocation and mood regulation during music listening, correlated positively with all BANSEQ facets, particularly with the BANSEQ *mood regulation through singing* facet (*r* = .53; *p* < .001), but was not linked to *social singing reward*. The *performativity* facet of MusEQ, reflecting daily music-making behaviours, also correlated positively with all BANSEQ facets, most strongly with the *singing frequency* facet (*r* = .39; *p* < .001).

**Figure 3.**
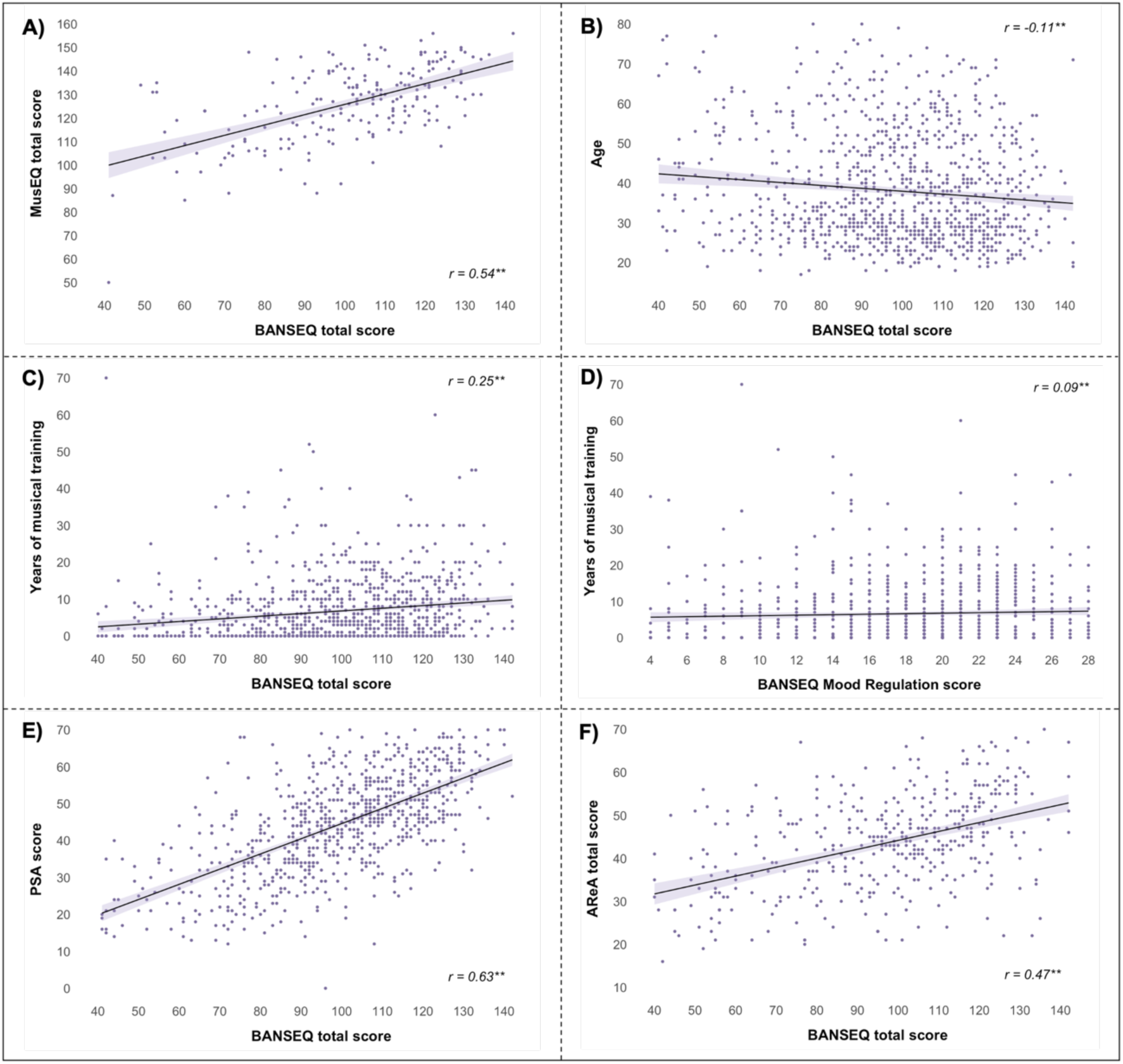
Correlation plots of BANSEQ scores with selected sociodemographic and psychological measures: (**A**) MusEQ total score with BANSEQ total score, (**B**) age with BANSEQ total score, (**C**) years of prior musical training with BANSEQ total score, (**D**) years of prior musical training with BANSEQ Mood Regulation facet, (**E**) PSA total score with BANSEQ total score, and (**F**) AReA total score with BANSEQ total score. Each point represents an individual participant; solid lines denote linear regression fits; and shaded bands indicate 95% confidence intervals. (BANSEQ: Barcelona-Aarhus Natural Singing Engagement Questionnaire; MusEQ: Music Engagement Questionnaire; PSA: Perceived Singing Abilities; AReA: Aesthetic Responsiveness Assessment). *\* p < .05; ** p < .001*.

**Table 7.**
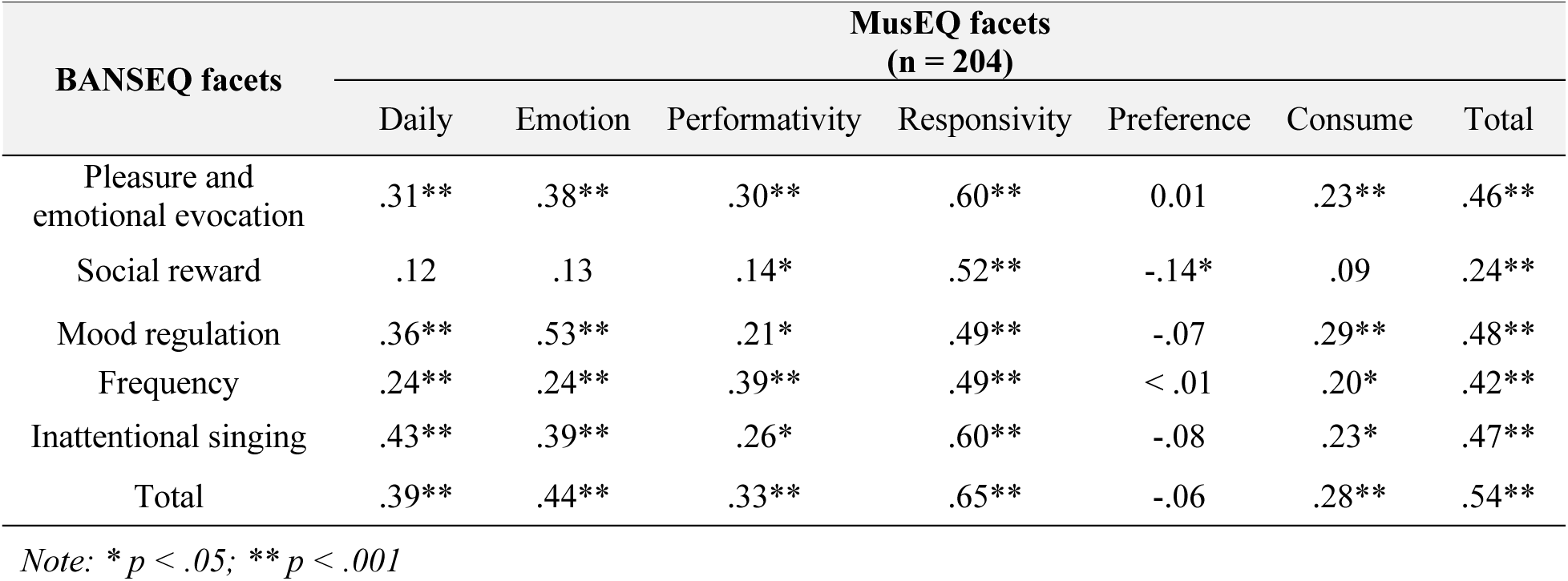
Correlations between the facets and total scores of the BANSEQ and the MusEQ. Spearman’s Rho (*r*) are presented per each comparison. (BANSEQ: Barcelona-Aarhus Natural Singing Engagement Questionnaire; MusEQ: Music Engagement Questionnaire).

| BANSEQ facets | MusEQ facets<br>(n = 204) |  |  |  |  |  |  |
| --- | --- | --- | --- | --- | --- | --- | --- |
|  | Daily | Emotion | Performativity | Responsivity | Preference | Consume | Total |
| Pleasure and emotional evocation | .31** | .38** | .30** | .60** | 0.01 | .23** | .46** |
| Social reward | .12 | .13 | .14* | .52** | -.14* | .09 | .24** |
| Mood regulation | .36** | .53** | .21* | .49** | -.07 | .29** | .48** |
| Frequency | .24** | .24** | .39** | .49** | < .01 | .20* | .42** |
| Inattentional singing | .43** | .39** | .26* | .60** | -.08 | .23* | .47** |
| Total | .39** | .44** | .33** | .65** | -.06 | .28** | .54** |

Note: \* $p < .05$ ; \*\* $p < .001$

#### Relationship of the BANSEQ with sociodemographic, musical and personality factors Effects of age and gender

Older age showed small negative correlations with the BANSEQ total score (*r* = -.11; *p* > .001), *inattentional singing* (*r* = -.17; *p <* .001), *mood regulation through singing* (*r* = -.10; *p <* .014), and *pleasure and emotional evocation* (*r* = -.16; *p <* .001) (see *Figure 3.B*), whereas no correlations were found between age with *singing frequency* or *social singing reward*.

When comparing levels of natural singing engagement across women, men, and non-binary participants, all BANSEQ total and facet scores differed by gender. *Post-hoc* comparisons with Bonferroni corrections revealed that all significant differences were observed between women and men. Women reported significantly higher scores than men in BANSEQ total (*H* (2) = 34.9, *p <* .001), *singing frequency* (*H* (2) = 25.7, *p <* .001), *social singing reward* (*H* (2) = 33, *p <* .001), *pleasure and emotional evocation* (*H* (2) = 57.5, *p <* .001), *mood regulation through singing* (*H* (2) = 29.6, *p <* .001), and *inattentional singing* (*H* (2) = 16.8, *p <* .001) scores.

#### Effects of prior musical experience and self-perceived singing abilities

Participants with more years of musical training showed higher BANSEQ total score (see *Figure 3.C*). The strongest association was with the BANSEQ *singing frequency* facet (*r* = .33; *p <* .001), while the weakest with *mood regulation through singing* (*r* = .09; *p* > .018) (see *Figure 3.D*). Similarly, higher PSA was positively correlated with all facets and the overall BANSEQ score (*r* = .63; *p* < .001) (see *Figure 3.E*). The strongest association was observed with *singing frequency* (*r* = .69; *p <* .001), while the weakest association with *mood regulation through singing* (*r* = .34; *p <* .001) (see *Table 8*).

**Table 8.** Correlations between the facets and total scores of the BANSEQ and the years of prior musical training and PSA (Perceived Singing Abilities). Spearman’s Rho (*r*) are presented per each comparison.

| BANSEQ facets | Years of musical training<br>(n = 1031) | Perceived Singing Abilities<br>(n = 810) |
| --- | --- | --- |
| Pleasure and emotional evocation | .24** | .58** |
| Social reward | .27** | .52** |
| Mood regulation | .09* | .34** |
| Frequency | .33** | .69** |
| Inattentional singing | .20** | .44** |
| Total | .25** | .63** |

Note: \* $p < .05$ ; \*\* $p < .001$

#### Effects of aesthetic responsiveness and other personality traits

To explore how natural singing engagement relates to broader dimensions of artistic experience and personality traits, the BANSEQ (total and facets scores) was correlated with the AReA (total and facet scores) and the BFI-2-XS (facet scores) (see *Table 9*). The BANSEQ total and all facet scores correlated positively with the AReA total and all facet scores. The strongest association for the BANSEQ overall score was with the AReA total score (*r* = .47; *p <* .001) (see *Figure 3.F*), followed by the AReA *aesthetic appreciation* facet (*r* = .45; *p <* .001). Indeed, the AReA *aesthetic appreciation* facet was the most strongly correlated with the BANSEQ facets of *mood regulation*, *inattentional singing*, *pleasure and emotional evocation*, and *social singing reward*, compared to other AReA facets.

**Table 9.** Correlations between the facets and total scores of the BANSEQ and the AReA and BFI-2-XS. Spearman’s Rho (*r*) are presented per each comparison. (BANSEQ: Barcelona-Aarhus Natural Singing Engagement Questionnaire; AReA: Aesthetic Responsiveness Assessment; BFI-2-XS: extra-short form of the Big Five Inventory-II; Extr.: Extraversion; Agree.: Agreeableness; Cons.: Conscientiousness; Neur.: Neuroticism; Open.: Openness).

| BANSEQ facets | AReA (n = 430) |  |  |  | BFI-2-XS (n = 204) |  |  |  |  |
| --- | --- | --- | --- | --- | --- | --- | --- | --- | --- |
|  | Appreciation | Experience | Creative | Total | Extr. | Agree. | Cons. | Neur. | Open. |
| Pleasure and emotional evocation | .37** | .32** | .31** | .39** | .15* | .10 | .05 | .05 | .16* |
| Social reward | .32** | .27** | .21** | .31** | .26** | .13 | .03 | -.07 | .09 |
| Mood regulation | .43** | .41** | .30** | .44** | .16* | .14* | .19 | .07 | .05 |
| Singing frequency | .37** | .37** | .35** | .41** | .14* | .06 | .05 | .13 | .14* |
| Inattentional singing | .40** | .37** | .30** | .41** | .14* | .09 | -.03 | .05 | .09 |
| Total | .45** | .42** | .35** | .47** | .18* | .12 | .05 | .06 | .13* |

Note: \* $p < .05$ ; \*\* $p < .001$

Regarding personality traits, the BANSEQ total and all facet scores correlated positively to *extraversion*, with the strongest association observed for the *social singing reward* facet (*r* = .26; *p <* .001), followed by *mood regulation through singing* facet (*r* = .16; *p <* .001). The trait *openness* showed the strongest correlation with the BANSEQ *pleasure and emotional evocation* (*r* = .16; *p <* .022), followed by *singing frequency* and BANSEQ total score. The trait *agreeableness* correlated only with *mood regulation through singing* (*r* = .14; *p <* .042), while no significant associations were found between any BANSEQ facet or total score and *conscientiousness* or *neuroticism*.

## Discussion

This research aimed to identify the underlying facets that explain variability of reward responses to natural singing in the general population. Natural singing is a form of music-making that does not require formal musical training and is expected to share motivational components with broader music reward and engagement processes. Given its lack of direct biological advantages, a central question is why humans derive reward from singing across cultures and throughout the lifespan, often in the absence of formal training, technical mastery, or performance goals. Building on prior research on individual differences in music reward and engagement, we developed an initial pool of items capturing the rewarding experiences of non-professional everyday singing to characterize the distinct facets underlying variability in the hedonic experience of singing. These items were administered to two international samples of adults with advanced or proficient English, and factorial analyses confirmed the five latent facets characterizing variability in sensitivity to natural singing reward: (1) *pleasure and emotional evocation*, (2) *social singing reward*, (3) *singing frequency*, (4) *mood regulation through singing*, and (5) *inattentional singing during routine tasks*. Although a unidimensional structure showed acceptable fit, the five-factor solution demonstrated excellent goodness-of-fit indices, with factors strongly intercorrelated, indicating that they capture related but distinguishable aspects of natural singing that account for variability in sensitivity to singing reward. Based on these findings, we developed a new self-report psychometric instrument called Barcelona-Aarhus Natural Singing Engagement Questionnaire (BANSEQ), which assesses sensitivity to natural singing reward in the general population across the five identified facets. The BANSEQ exhibited excellent reliability (α = .941), and proved to be stable and replicable at the population level.

Examining the extent to which natural singing enjoyment overlaps with established measures of music reward and engagement provides insight into the shared aspects that drive intrinsic motivation for singing, music listening, and other forms of music-making. For this reason, we explored the association of BANSEQ facets with two validated scales assessing music reward (eBMRQ) and music engagement (MusEQ), both of which demonstrate high reliability (α = .92-.95). The eBMRQ captures individual differences in reward derived from passive music exposure across six facets, including musical seeking, emotional evocation, mood regulation, social reward, sensory-motor responses, and absorption (Cardona et al., 2022; Mas-Herrero et al., 2013). The MusEQ assesses individual differences in daily music engagement — closely related to music reward — through facets such as everyday listening, emotional and mood-related responses to music, engagement in music-making behaviours, responsiveness to music via inattentive, social, or individual singing and movement, breadth of musical taste, and music consumption habits (Vanstone et al., 2016). Furthermore, we examined which sociodemographic and psychological factors are associated to each facet underlying variability in singing reward. In the following subsections, we discuss the facets that characterize sensitivity to natural singing reward, and the individual factors associated with each.

### Pleasure and emotional evocation

This facet captures the consummatory pleasure and emotions elicited by singing itself, independent of external or contextual factors. Given the key role of emotional processing in the pleasure derived from music listening (Vuilleumier & Trost, 2015), it is unsurprising that pleasure and affective states overlap in singing reward. The fact that humans often enjoy singing in solitary contexts purely for pleasure, without explicit social or mastery goals (e.g., singing in the shower) highlights its direct engagement with the reward system. Neuroimaging studies have shown that pleasurable music activates the ventral striatum, including the nucleus accumbens, and increases dopamine release in this region, which is also responsive to other rewarding stimuli and to music evoking joy and happiness (Koelsch et al., 2006; Mitterschiffthaler et al., 2007; Salimpoor et al., 2011). Beyond basic affective categories with clear adaptive functions (e.g., joy or sadness), music can evoke a wide range of complex or aesthetic emotions that often arise from cognitive associations or autobiographical memories, such as wonder, tenderness, nostalgia, tension, power, or peacefulness (Strauss et al., 2024). While complex music-derived emotions with positive valence have been associated with increased activity in ventral striatum and ventral tegmental area, negatively valanced complex emotions have been linked to greater orbitofrontal cortex activity, showing consistency with broader models of emotion processing (Vuilleumier & Trost, 2015). In this vein, singing may engage similar neural circuits involved in pleasure and emotional processing. Enjoyment of solitary singing may also be closely related to other BANSEQ facets, such as the use of singing for mood regulation.

### Social singing reward

Although singing can be intrinsically pleasurable in solitary contexts, it often occurs in culturally embedded social settings. Beyond familiar occasions in which people sing together, such as birthdays or religious ceremonies, choral singing also constitutes a meaningful leisure activity for many individuals. Theories of the evolutionary significance of singing propose functions related to strengthening social bonds or serving as a byproduct of adaptations for emotional expression and communication (Bryant, 2013; Launay & Pearce, 2020; Mehr et al., 2021; Savage et al., 2021). Given that belongingness is a fundamental psychological need essential for humans’ social development, well-being, and optimal functioning (Mellor et al., 2008), singing may serve as a behaviour that enhances emotional expression, shared affect, and affiliation with others. In this context, the rewarding value of singing may be amplified by interpersonal interaction and increased group cohesion. Singing provides a unique opportunity to use the voice collectively to produce a shared output with aesthetic, communicative, or ritualistic functions, with its emotional impact experienced jointly by group members (Keeler et al., 2015). Singing in unison requires real-time synchronization of rhythm, timing, pitch, and melody, which in turn promotes alignment of respiratory, cardiac, and sensorimotor processes across individuals (Delius & Müller, 2023; Pearce et al., 2016). Such interpersonal synchrony has been shown to enhance prosocial behaviours (e.g., helping, sharing, trusting) and prosocial dispositions (e.g., affiliation, empathy, perceived closeness, sense of community), and it is associated with increased endorphin release, contributing to more positive social evaluations among group members (Hu et al., 2022; Pearce et al., 2015). The cooperative demands of group singing, including attentive listening and mutual performance adjustment, are key mechanisms supporting social bonding. Moreover, the shared achievement of producing a cohesive vocal output may serve as a marker of successful coordination, reinforcing feelings of integration and motivation to continue cooperating (Pearce et al., 2017). Shared strong emotional experiences during singing further contribute to group cohesion. For example, national or team anthems are often designed to be emotionally arousing, which underscores the role of shared affect in fostering collective identity (Juslin & Västfjäll, 2008). Neuroimaging studies have demonstrated brain-to-brain synchronization during singing or humming, particularly in frontal regions involved in motor, language, and executive functions, with such synchronization correlating with coordinated performance and social bonding (Delius & Müller, 2023). Consistent with this, choral singing has shown to enhance feelings of belongingness, trust, cooperation, and community involvement more than other group artistic activities, such as poetry reading, film viewing, or passive music listening (Keeler et al., 2015; Pearce et al., 2016). The *social singing reward* facet of the BANSEQ captures a distinct dimension of natural singing engagement, demonstrating the variability in the extent to which individuals experience singing as socially rewarding.

### Mood regulation through singing

Singing provokes multiple physiological changes. Beyond effects on the cardiorespiratory system, singing can upregulate neurotransmitters and hormones linked to positive affect and downregulate stress-related hormones (Kang et al., 2017). This behaviour requires volitional breath control, precise laryngeal muscle coordination, and dynamic shaping of the vocal tract to achieve the desired sound. Continuous awareness of airflow, muscle engagement, and articulator configuration enables singers to optimize acoustic output and expressive control (Wild, 2004). This ongoing monitoring and prediction of bodily states reflects interoception, an adaptive brain process fundamental to emotion processing that has been linked to a range of affective states, including joy, anger, and aversion (Feldman et al., 2024). Therefore, singing with emotional intent relies on precise perception and regulation of internal physiological signals, making singing a deeply embodied activity (Kleber et al., 2013). The embodied multisensory integration of personally relevant internal and external cues has been suggested to enhance awareness of bodily sensations and, in turn, facilitate their modulation (Ceunen et al., 2016). In this sense, singing may function as an intentional but implicit activity grounded in the body’s innate regulatory capacities and shaped by lyrical content and emotional intent, leading to mood regulation (Juntunen et al., 2023; Peters et al., 2024). In a daily basis, individuals listen to music to shape mood states and modulate emotional arousal (Baltazar & Saarikallio, 2016). Prior neuroimaging studies have shown that musical experiences engage neural circuits implicated in emotion processing, supporting the notion that music engagement facilitates emotional regulation (Peters et al., 2024). Mood regulation through singing may thus operate through mechanisms similar to those involved in music listening, while being further potentiated by the enhanced emotional expression and embodied engagement inherent to vocal performance. Although the precise mechanisms underlying mood regulation through singing remain to be fully elucidated, both individual and group singing have been shown to enhance mood, increase positive affect, and reduce stress in the general population (Fancourt et al., 2015). The BANSEQ captures this facet of natural singing engagement as a distinct pathway through which singing becomes rewarding. Changes in mood during singing might be influenced by both emotional evocation and consummatory pleasure, leading to heightened positive affective states. For some individuals, participation in social singing contexts may further enhance mood through additional rewards derived from shared musical engagement.

### Singing frequency

An important indicator of intrinsic motivation to engage in any activity is the amount of time individuals voluntarily devote to it (Deci & Ryan, 2000). Given the high accessibility of singing in humans, who can sing at many moments throughout the day without being heard (e.g., in the shower or while driving), the frequency with which individuals engage in natural singing may directly reflect sensitivity to singing reward and engagement. However, particularly in adulthood, engagement in singing can be constrained by contextual factors such as work demands or caregiving responsibilities, which may reduce opportunities for musical activity (Theorell et al., 2015). Despite the ubiquity of singing across cultures, participation rates vary across countries and across the lifespan (Shilton et al., 2023). A prior study with population-level data from Sweden indicated that engagement in musical activities, including singing, decreases between the ages of 27 and 54, reflecting the tendency for many individuals to discontinue these activities in adulthood. In the study, around 10% of teenagers reported singing in a choir at least once per week, compared to 5% of young adults (Theorell et al., 2015). Although contextual or personal constraints can limit singing engagement, the low barriers and high availability of singing allow the singing frequency facet of the BANSEQ to capture a behavioural manifestation of sensitivity to singing reward and, consequently, individual differences in intrinsic motivation to engage in singing within the general population.

### Inattentional singing during routine tasks

In everyday life, many activities are performed routinely and with minimal cognitive demand, such as cleaning or walking. During such low-load tasks, some individuals experience what we term *inattentional singing*, a facet of natural singing behaviour that we have identified as positively associated with overall singing reward. This phenomenon overlaps with prior descriptions of inattentive musical imagery, commonly referred to as earworms, which are experiences in which fragments of music are spontaneously and repetitively activated in the mind without a clear sense of agency (Beaman & Williams, 2010). Large-scale survey data indicate that earworms are highly prevalent, with nearly 90% of individuals reporting their occurrence on a weekly basis. Although they are involuntary, the music involved is typically familiar and well liked, suggesting a link to musical reward (Williamson et al., 2014). Some theories propose that earworms are more likely to involve songs that are easy to sing, supporting a close relationship between involuntary musical imagery and singing behaviour (Killingly et al., 2021; Theorell et al., 2019). Earworms have been associated with subjective singing ability and the tendency to sing along with music in everyday life. Indeed, evidence suggests that singing aloud increases the likelihood of experiencing earworms, while a common response to having a song stuck in one’s head is to sing (Müllensiefen et al., 2014). From this perspective, inattentional singing can be understood as a form of automatic inner singing. Moreover, inattentional singing may also serve as tool for mood regulation. Since music has been shown to distract individuals from daily worries and to interrupt maladaptive thought patterns (Theorell et al., 2019), inattentional singing of a familiar or pleasurable song may function as an unconscious regulatory mechanism, filling moments of cognitive underload or providing a rhythmic structure during repetitive activities, such as walking or upper-limb movements in household chores. Given that earworms and inattentional singing are more prevalent among individuals with higher levels of singing engagement, it is unsurprising that this facet emerged as a distinct facet of the BANSEQ, contributing to the modulation of natural singing reward in the general population.

### The distinct profiles of singing and musical hedonism

Although singing is a form of music-making, the present study demonstrates that natural singing reward, as measured by the BANSEQ, is only partially overlapping with broader music-related reward and engagement. Specifically, singing reward shared 34% of its variance with music reward (eBMRQ) and 30% with music engagement (MusEQ) in the general population. When examining convergence with sensitivity to reward across other sensory domains (measured by the SHAPS), smaller but still meaningful correlations indicated that general physical and social anhedonia are also associated with lower levels of singing reward, consistent with previous findings on musical anhedonia (Mas-Herrero et al., 2013). These patterns indicate that whereas natural singing engagement is related to music-specific reward processes more than to general hedonic capacity, it still retains substantial variance not explained by passive exposure to music alone or other forms of music engagement. Facet-level analyses revealed that the strongest associations between singing and music reward and engagement was *social reward* and *mood regulation*. Similarly, the *mood regulation* facet showed strong convergence between singing reward and music engagement, while the *responsivity* facet of music engagement (encompassing inattentive, social, and individual singing) was strongly associated with *pleasure and emotional evocation* during singing. These findings suggest that certain motivational mechanisms, particularly those related to social and affective regulation, are shared across singing and music reward, even when overall hedonic responses to music differ. At the same time, the partial overlap and differential strength of facet-level associations indicate that intrinsic motivation to sing cannot be fully reduced to enjoyment of music as an auditory stimulus, pointing instead to additional vocal- and action-based sources of reward.

By examining of the joint distribution of BANSEQ and eBMRQ scores, we observed distinct profiles of singing and musical hedonism, reflecting individual differences in the capacity to derive reward from singing versus music listening. After classifying participants as higher, normal, and lower hedonic for singing and music (based on overall scores falling above, below, or within ±1.5 SD of the mean on both scales), 77.55% of them fell within the normal range for both forms of hedonism (see *Figure 1*). Notably, 3.56% of individuals reported a preference for engaging in singing over passive music exposure (P-Sing), showing normal levels of singing reward but reduced music reward. For these individuals, the rewarding experience of singing may not be primarily driven by auditory perception or musical consumption, but rather by the act of vocal production itself, constituting a partially independent source of reward. Conversely, 2.92% of participants showed normal sensitivity to music reward but reduced singing reward (P-Music), indicating that enjoyment of music listening does not necessarily translate into motivation to sing. This finding suggests that singing may be experienced as effortful, socially inhibiting, evaluative, or non-aesthetic for some individuals, requiring additional motivational components beyond perceptual musical pleasure. To further characterize these distinct profiles, we examined BANSEQ facet scores within the P-Sing and P-Music groups (see *Figure 2* and *Table 6*). For individuals who preferred music over singing (P-Music), singing reward was primarily driven by *pleasure and emotional evocation* and by *social singing reward*, whereas *singing frequency* showed the lowest contribution. In contrast, individuals who preferred singing over music (P-Sing) displayed a similar emphasis on pleasure and social reward, but also showed elevated levels of *inattentional singing*, reaching values comparable to those of the full sample. Among participants with low overall singing enjoyment (i.e., singing anhedonia), *social singing* emerged as the facet through which they experienced the greatest relative enjoyment, followed by *pleasure and emotional evocation*. Conversely, individuals with the highest sensitivity to singing reward (i.e., singing hyperhedonia) exhibited elevated scores across all facets, but particularly in *pleasure and emotional evocation* and *inattentional singing*. Taken together, these patterns suggest a graded structure of singing reward in the general population. Social interaction-related reward may constitute the primary entry point for singing engagement, particularly among individuals with low overall singing enjoyment. Pleasure and emotional evocation, along with inattentional singing, appear to support typical levels of singing reward, while the use of singing for mood regulation and higher singing frequency characterize individuals at the upper end of the singing hedonism spectrum. This progression highlights the multifaceted nature of singing reward and supports the view that natural singing constitutes a distinct domain of human hedonic experience.

### Factors modulating sensitivity to natural singing reward

The extent to which humans experience musical activities as rewarding can be shaped by multiple individual characteristics (Mas-Herrero et al., 2013; Tang, 2026). Despite singing is present across the human lifespan, the present sample (aged 18-80 years) revealed that older participants reported slightly lower overall singing reward, particularly in *inattentional singing*, *mood regulation*, and *pleasure and emotional evocation*. Although these associations are modest, they are in line with prior findings in the music reward literature where increasing age was linked to reduced mood regulation through music (Cardona et al., 2022; Mas-Herrero et al., 2013). This finding is consistent with evidence that healthy aging is accompanied by functional alterations in the dopaminergic reward system (Dreher et al., 2008). Notably, age was not associated with *social singing reward* or *singing frequency*, possibly reflecting the enduring presence of singing in social and cultural contexts across the lifespan, such as religious ceremonies and traditional celebrations, in which singing remains common regardless of age (Trehub & Trainor, 1998).

Gender differences also emerged in natural singing reward, with women reporting higher levels across all BANSEQ facets. This finding aligns with prior evidence showing greater sensitivity to music reward in women (Mas-Herrero et al., 2013). From an evolutionary perspective, music-making behaviours have been proposed to play a role in caregiving and early bonding, particularly in the context of parent-infant interactions (Theorell et al., 2015). Singing during pregnancy and early infancy has been shown to influence infant emotional responses and stress regulation more strongly than passive music listening, while also enhancing caregivers’ feelings of closeness, calmness, and emotional stability (Markova et al., 2020). The tendency of some caregivers to sing lullabies or play-songs has also been shown to be particularly effective in helping infants sleep or increasing their attention in interactions, respectively (Cirelli et al., 2019). Although caregiving roles are increasingly attributed to people of all genders in contemporary societies, the higher biological investment of energy of women in reproduction (i.e., pregnancy and newborn care) may partly explain their heightened sensitivity to singing reward and engagement (Walzer, 1996). An alternative adaptationist theory has proposed that music evolved as a signalling system for intelligence, particularly in mating contexts (Miller, 2000). Some evidence suggests that musical production abilities may enhance perceived mate value, especially among less physically attractive males. However, musical performance quality does not clearly indicate health status or parenting ability, suggesting that it may function as a broad indicator of cognitive engagement rather than a specialized mating adaptation (Madison et al., 2018), which could explain why men did not showed higher sensitivity to singing reward compared to women.

Greater years of musical training were associated with higher overall singing reward, and particularly with *singing frequency*, an expected result given that any type of musical training is often supported by internal singing (Casas-Mas et al., 2019). In line with prior research showing that mood regulation through music listening does not differ as a function of musical expertise (Mas-Herrero et al., 2013), musical training was not associated with *mood regulation* through singing. A similar pattern emerged for self-perceived singing ability: although individuals who rated their skills higher reported singing more frequently, the association with *mood regulation* was comparatively weak. These results suggest that emotional benefits derived from natural singing do not require high levels of vocal mastery nor prior formal training, supporting the idea that singing may have adaptive value for emotional well-being among the general population.

Individual differences in aesthetic responsiveness were also associated with overall sensitivity to singing reward, with the strongest association observed for *mood regulation* through singing. While reward derived from music listening has been linked to higher aesthetic responsiveness (Mas-Herrero et al., 2013), singing may provide an additional reward value when the vocal act produces an aesthetically satisfying auditory outcome. Finally, personality traits also modulated sensitivity to singing reward. Overall singing reward was positively associated with extraversion, particularly through *social singing reward*, which is in line with prior findings linking social aspects of musical hedonism to extraversion (Wang et al., 2021). Openness showed the strongest association with *pleasure and emotional evocation* during singing, while agreeableness was most strongly related to *mood regulation* through singing. These patterns align with prior research linking music reward to openness and agreeableness, while contrasting with their findings on the influence of neuroticism and conscientiousness (Theorell et al., 2015; Wang et al., 2021). The expressive and performative nature of singing may therefore be especially appealing to individuals who are social, cooperative, and creative, while being less enjoyed by highly neurotic or conscientious individuals, who are more prone to negative affect and may prefer more structured and controllable activities (Soto & John, 2017).

### Strengths, limitations, and future directions

The present research identified the latent facets that explain variability in sensitivity to natural singing reward among the general population. Notably, the items used to measure the hedonic experience of singing were designed independently of individuals’ musical experience, singing skills, and intercultural variability in singing practices. For this reason, we recruited an international adult sample encompassing ten nationality clusters based on the Global Clustering of Countries by Culture (Mensah & Chen, 2012). Although some societies practice group singing in a greater extent than others, recent research has suggested that between-region differences in personality profiles are not associated with music reward. This indicates that music-related reward processes are broadly comparable across geographical regions, with within-region variation in self-construal better explaining individual differences (Tang, 2026). This research led to the development of the BANSEQ, a new psychometric tool that assess reward sensitivity to natural singing in the general population and that was validated in an international sample. However, future studies should further examine its ecological validity in more culturally balanced samples, as 68.1% of participants in the present research were from Anglo-Saxon backgrounds. A further strength of the present research is the wide age range of participants (18-80 years), allowing examination of singing reward across the adult lifespan. However, the sample was skewed toward younger adults (mean age = 38), with only 8.8% of participants aged 60-80. This underrepresentation of older adults may limit generalizability, particularly given that older individuals may have greater time availability for leisure activities and community-based singing, which could influence engagement patterns. Future work should therefore recruit larger and more representative samples of older adults.

Having shown that variability in sensitivity to singing reward can be explained by facets involving emotional processing, future research includes the use of behavioural assessments to investigate the complex emotions elicited by singing. Further research should also incorporate psychophysiological and neuroimaging methods to explore the neural and bodily mechanisms underlying singing reward. Given the reported benefits of singing for both physical and mental health in clinical and non-clinical populations, the BANSEQ may also be useful in guiding the design of singing-based interventions aimed at supporting neurological and psychological functions associated with reward processing. By assessing individuals’ levels of singing engagement, such interventions could potentially be tailored to maximize benefits and to help predict their effectiveness.

## Conclusions

The present research demonstrates that individuals differ in their capacity to derive reward from natural (non-professional) singing within the general population, regardless of prior training or skills. Natural singing reward showed to be modulated by five facets: *pleasure and emotional evocation*, *social singing reward*, *mood regulation through singing*, *singing frequency*, and *inattentional singing during routine tasks*. Although singing reward partially overlaps with broader music reward processes, the present findings show that the performative act of singing entails distinct reward-related mechanisms, particularly within social and affective domains. Individual differences in sociodemographic and personal characteristics such as age, gender, self-perceived singing skills and personality traits further modulate sensitivity to singing reward across these facets. As a result of the present research, we developed the Barcelona-Aarhus Natural Singing Engagement Questionnaire (BANSEQ), a psychometric tool that assesses reward sensitivity to natural singing in the general population across five facets underlying the hedonic capacity of singing in humans.

## Acknowledgements

We thank the participants who voluntarily completed the surveys, as well as Xim Cerda-Company, Silvia Rondini, and Samantha Vigo for their assistance with data collection. This research was funded by Ministerio de Ciencia e Innovación, AEI /10.13039/501100011033/ and ERDF - A way of making Europe, as well as by the European Union. Views and opinions expressed are however those of the author(s) only and do not necessarily reflect those of the European Union or the European Health and Digital Executive Agency (HaDEA). Neither the European Union nor the granting authority can be held responsible for them. We also acknowledge institutional support from the CERCA Programme / Generalitat de Catalunya. ARF was supported by the FIAS Fellowship Programme, and co-funded by the European Commission through the Marie Skłodowska-Curie Actions COFUND programme (Grant No. 945408). BK was supported by the Carlsberg Foundation (Grant No. CF22-1172).

## Appendix

### A. The Barcelona-Aarhus Natural Singing Engagement Questionnaire

The following questionnaire is about the extent that you might like singing or engaging in activities related to singing. Importantly, this is not a questionnaire that pretends to measure singing as a professional activity or being trained as a singer or the extent that you are a very good singer. Indeed, we are conducting this research in order to understand better natural singing behavior, which appears in infancy and represent an important musical ability for humans.

Each item is a statement with which you can agree or disagree. For each statement, please indicate the extent to which you agree or disagree. Please answer all items, do not leave any blank. Choose only one answer for each statement. Please try to be as accurate and honest as possible, answering each point as if it were the only one. Do not worry about being “consistent” in your answers. As always in this type of questionnaires, there are not correct answers, we only aim to investigate individual differences in natural singing. Choose for each item the following options:

[1] - Strongly disagree; [2] - Disagree; [3] - Somewhat disagree; [4] Neither agree nor disagree; [5] - Somewhat agree; [6] - Agree; [7] - Strongly agree
1. I sing weekly alone or in a group.
2. I become cheerful when I sing in a group.
3. I enjoy singing.
4. When I am sad, I sing certain songs to feel better.
5. I sing or hum as I go about my daily activities.
6. I have practiced singing on a regular basis during my lifetime.
7. If others are singing, I join in.
8. I sing by myself for pleasure.
9. Singing takes me back to my teen years.
10. I sing or hum when doing certain tasks (e.g., cooking or cleaning).
11. There are times when I do not nothing except singing.
12. I feel uncomfortable when signing in a group.
13. I feel happy when singing by myself.
14. When someone hurts my feelings, I sing songs to deal with it.
15. I sing or hum while hiking or travelling.
16. I do sing or practice singing for a certain amount of time per day.
17. I hate social activities related to singing (e.g., going to karaoke).
18. I learn to sing songs that I love regardless of whether I will show it to someone else or not.
19. I become emotional when I sing certain songs.
20. If I am bored, I sing or hum a song to pass the time (e.g., performing chores, waiting in a queue, etc.)

