## Supplementary Material for "Characterizing reward sensitivity to natural singing: an individual differences approach"

[h] Institució Catalana de Recerca i Estudis Avançats, Barcelona, Spain.

* Both authors declare **co-senior equal contribution**

**Corresponding author:**

Antoni Rodríguez-Fornells

ICREA Research Professor

Department of Cognition, Development and Educational Psychology

University of Barcelona

Campus Bellvitge, Feixa Llarga, s/n, 08907

L'Hospitalet de Llobregat, 08907 Barcelona, Spain

### Supplementary material

**Supplementary Table 1**. Participants' nationality of the total sample

| Nationality | % |
|---|---|
| American | 20.4 |
| Spanish | 17.0 |
| British | 13.2 |
| French | 7.8 |
| Canadian | 6.7 |
| Danish | 4.2 |
| Australian | 4.1 |
| Italian | 3.4 |
| Colombian | 2.0 |
| German | 1.9 |
| Greek | 1.3 |
| Chinese | 1.2 |
| Dutch | 1.2 |
| Indian | 1.0 |
| Brazilian | 1.0 |
| Mexican | 0.9 |
| Cuban | 0.8 |
| Finnish | 0.7 |
| Irish | 0.7 |
| Belgian | 0.6 |
| Chilean | 0.6 |
| Polish | 0.6 |
| Others | 9.1 |

**Supplementary Table 2**. Factor loadings for the unidimensional solution and item loadings across facets in Study 1 (n = 606; exploratory factor analysis sample). Salient loadings are printed in bold. (BANSEQ: Barcelona-Aarhus Natural Singing Engagement Questionnaire; SF: Singing Frequency; SR: Social Reward; PE: Pleasure and Emotional evocation; MR: Mood Regulation; IS: Inattentional Singing).

| Item | | BANSEQ | SF | SR | PE | MR | IS |
|---|---|---|---|---|---|---|---|
| 1 | *I sing weekly alone or in a group.* | **.628** | **.533** | .118 | .079 | -.066 | .122 |
| 2 | *I become cheerful when I sing in a group.* | **.527** | .023 | **.713** | .072 | -.040 | .027 |
| 3 | *I enjoy singing.* | **.689** | .250 | .119 | **.696** | -.164 | -.066 |
| 4 | *When I am sad, I sing certain songs to feel better.* | **.578** | -.074 | -.012 | -.023 | **.909** | .033 |
| 5 | *I sing or hum as I go about my daily activities.* | **.693** | -.018 | .001 | -.022 | -.014 | **.925** |
| 6 | *I have practiced singing on a regular basis during my lifetime.* | **.563** | **.809** | .042 | -.022 | -.074 | -.038 |
| 7 | *If others are singing, I join in.* | **.553** | -.049 | **.708** | -.013 | .037 | .151 |
| 8 | *I sing by myself for pleasure.* | **.753** | .086 | -.069 | **.713** | .066 | .097 |
| 9 | *Singing takes me back to my teen years.* | **.409** | -.047 | .141 | .018 | **.399** | .055 |
| 10 | *I sing or hum when doing certain tasks (e.g., cooking or cleaning).* | **.589** | -.068 | -.062 | .134 | -.071 | **.775** |
| 11 | *There are times when I do not nothing except singing.* | **.587** | **.607** | -.066 | -.040 | .094 | .134 |
| 12 | *I feel uncomfortable when signing in a group.* | **-.397** | -.143 | **-.706** | .064 | .067 | .081 |
| 13 | *I feel happy when singing by myself.* | **.682** | -.029 | -.042 | **.812** | .033 | .036 |
| 14 | When someone hurts my feelings, I sing songs to deal with it. | **.570** | .166 | -.056 | .038 | **.678** | -.054 |
| 15 | *I sing or hum while hiking or travelling.* | **.612** | .020 | .081 | -.013 | -.018 | **.698** |
| 16 | *I do sing or practice singing for a certain amount of time per day.* | **.663** | **.746** | .002 | .083 | .020 | -.020 |
| 17 | *I hate social activities related to singing (e.g., going to karaoke).* | **-.369** | .084 | **-.586** | -.019 | -.146 | .081 |
| 18 | *I learn to sing songs that I love regardless of whether I will show it to someone else or not.* | **.653** | .287 | -.032 | **.422** | .130 | -.019 |
| 19 | *I become emotional when I sing certain songs.* | **.563** | -.044 | .063 | .388 | **.432** | -.104 |
| 20 | *If I am bored, I sing or hum a song to pass the time (e.g., performing chores, waiting in a queue, etc.)* | **.656** | .094 | .019 | -.095 | .069 | **.739** |

**Supplementary Table 3**. Inter-factor correlation matrix for Study 1 (n = 606; exploratory factor analysis sample). (SF: Singing Frequency; SR: Social Reward; PE: Pleasure and Emotional evocation; MR: Mood Regulation; IS: Inattentional Singing).

| | SF | SR | PE | MR |
|---|---|---|---|---|
| **SR** | .562 | | | |
| **PE** | .365 | .484 | | |
| **MR** | .576 | .551 | .346 | |
| **IS** | .630 | .547 | .546 | .606 |

**Supplementary Table 4.** Factor loadings for the unidimensional solution and item loadings across facets in Study 2 (n = 430; confirmatory factor analysis sample). Salient loadings are printed in bold. (BANSEQ: Barcelona-Aarhus Natural Singing Engagement Questionnaire; SF: Singing Frequency; SR: Social Reward; PE: Pleasure and Emotional evocation; MR: Mood Regulation; IS: Inattentional Singing).

| Item | | BANSEQ | SF | SR | PE | MR | IS |
|---|---|---|---|---|---|---|---|
| 1 | *I sing weekly alone or in a group.* | **.690** | **.515** | 0 | 0 | 0 | 0 |
| 2 | *I become cheerful when I sing in a group.* | **.622** | 0 | **.620** | 0 | 0 | 0 |
| 3 | *I enjoy singing.* | **.835** | 0 | 0 | **.852** | 0 | 0 |
| 4 | *When I am sad, I sing certain songs to feel better.* | **.736** | 0 | 0 | 0 | **.750** | 0 |
| 5 | *I sing or hum as I go about my daily activities.* | **.719** | 0 | 0 | 0 | 0 | **.770** |
| 6 | *I have practiced singing on a regular basis during my lifetime.* | **.709** | **.832** | 0 | 0 | 0 | 0 |
| 7 | *If others are singing, I join in.* | **.668** | 0 | **.544** | 0 | 0 | 0 |
| 8 | *I sing by myself for pleasure.* | **.835** | 0 | 0 | **.679** | 0 | 0 |
| 9 | *Singing takes me back to my teen years.* | **.615** | 0 | 0 | 0 | **.753** | 0 |
| 10 | *I sing or hum when doing certain tasks (e.g., cooking or cleaning).* | **.685** | 0 | 0 | 0 | 0 | **.873** |
| 11 | *There are times when I do not nothing except singing.* | **.727** | **.506** | 0 | 0 | 0 | 0 |
| 12 | *I feel uncomfortable when signing in a group.* | **-.397** | 0 | **-.865** | 0 | 0 | 0 |
| 13 | *I feel happy when singing by myself.* | **.774** | 0 | 0 | **.687** | 0 | 0 |
| 14 | When someone hurts my feelings, I sing songs to deal with it. | **.667** | 0 | 0 | 0 | **.734** | 0 |
| 15 | *I sing or hum while hiking or travelling.* | **.658** | 0 | 0 | 0 | 0 | **.710** |
| 16 | *I do sing or practice singing for a certain amount of time per day.* | **.600** | **.875** | 0 | 0 | 0 | 0 |
| 17 | *I hate social activities related to singing (e.g., going to karaoke).* | **-.423** | 0 | **-.644** | 0 | 0 | 0 |
| 18 | *I learn to sing songs that I love regardless of whether I will show it to someone else or not.* | **.768** | 0 | 0 | **.511** | 0 | 0 |
| 19 | *I become emotional when I sing certain songs.* | **.722** | 0 | 0 | 0 | **.452** | 0 |
| 20 | *If I am bored, I sing or hum a song to pass the time (e.g., performing chores, waiting in a queue, etc.)* | **.693** | 0 | 0 | 0 | 0 | **.686** |

**Supplementary Table 5**. Inter-factor correlation matrix for Study 2 (n = 430; confirmatory factor analysis sample). (SF: Singing Frequency; SR: Social Reward; PE: Pleasure and Emotional evocation; MR: Mood Regulation; IS: Inattentional Singing).

| | SF | SR | PE | MR |
|---|---|---|---|---|
| **SR** | .591 | | | |
| **PE** | .679 | .652 | | |
| **MR** | .746 | .583 | .841 | |
| **IS** | .643 | .488 | .758 | .770 |
